# The Company Canids Confront: Spatiotemporal Partitioning at Local Scales Facilitates Carnivore Coexistence at the Landscape Level

**DOI:** 10.1101/2020.11.29.402529

**Authors:** Kadambari Devarajan, Abi Tamim Vanak

## Abstract

Canids are the most widely distributed carnivores in the world. The increasing impacts of commensal carnivores such as free-ranging dogs on wildlife communities has resulted in an urgent need to understand putative interactions within carnivore guilds. It is thus imperative to understand the processes driving canid assemblages in different landscapes and at multiple spatial and temporal scales, in order to conserve and manage wildlife communities. Here, we demonstrate the complex interactions and spatiotemporal dynamics underlying the coexistence of co-occurring carnivores in a landscape modified by the invasive, *Prosopis juliflora*. We investigated spatial, temporal, and habitat partitioning within a guild of four co-occurring canids in the arid northwest of India. The results indicate complex associations between the study species, where co-occurrence at the local spatial scale between species corresponds with temporal partitioning. Our study offers evidence that avoidance at the local scale and coexistence at the landscape scale are maintained in co-occurring intra-guild carnivores that are of similar body size through both facultative and behavioral character displacement such as temporal partitioning. It has also resulted in essential baseline information on the occurrence and distribution patterns of multiple canids in a human-dominated and understudied landscape threatened by global change. Understanding these biotic and abiotic drivers that impact carnivore guilds is crucial for the conservation and management of communities at the landscape scale.

## Introduction

Carnivores are distributed widely, found in almost all landscapes and land cover types on earth, and exhibit enormous variation in terms of traits and adaptations [Gittleman, 2013]. They influence community assembly and ecosystem function through their direct and indirect interactions with other species at different trophic levels, by exerting top-down control on a variety of prey species and affect that habitat use of other species through competitive interactions as well as intra-guild predation [Linnell & Strand, 2000; Gittleman, et al., 2001; Linnell, et al., 2001; Schuette, et al., 2013; Gompper, et al., 2016; Jiménez, et al., 2017].

Many vertebrate carnivore species are under enormous pressure from global change and large carnivores are thought to be particularly vulnerable to extinction risk from low population densities, inbreeding, diseases, climate change, habitat loss, poaching, and prey depletion [Linnell & Strand, 2000; Gittleman, et al., 2001; Linnell, et al., 2001; Schuette, et al., 2013]. Carnivores in resource-limited areas, such as arid and semi-arid ecosystems, are particularly vulnerable [Cardillo, et al., 2005; Ripple, et al., 2014].

While arid and semi-arid landscapes comprise a small fraction of the earth’s land surface, they harbor flora and fauna that are uniquely adapted to such landscapes and are often endemic to arid lands. However, these unique landscapes are threatened by encroachment, monocultures, invasive species, fragmentation, and climate change [Sodhi, et al., 2004]. Furthermore, little is known about the impact of species declines or extinctions on species interactions, while co-extinctions are increasing alarmingly in the anthropocene [Sodhi, et al., 2004; Dunn, et al., 2009]. As a consequence of human induced habitat loss, competing species are forced into smaller areas, thereby increasing the frequency of negative interactions. In countries such as India, savannas are under-studied, under-protected, and face rapid conversion [Vanak et al., 2016]. Despite these issues, species-rich carnivore communities continue to persist in heavily human-dominated and human-modified habitats.

Given the importance of carnivores, grasslands, and arid-lands, it is of high conservation priority to understand the processes and patterns driving carnivore community assemblages in resource-limited and depauperate landscapes. It is essential to understand whether and how species interactions influence their distribution patterns by testing for any apparent spatial, temporal, or habitat partitioning between co-occurring species. However, exploring such interactions is challenging since few places in the world support multiple sympatric carnivores belonging to the same guild.

Wild canids are the most widely distributed of carnivores and are found in all continents with the exception of Antarctica [Sillero-Zubiri, et al., 2004; Gittleman, 2013]. A common trend is that two or three species of canids tend to occur in sympatry [Kamler, et al., 2004; Gittleman, 2013; Kamler, et al., 2012]. Typically, in systems with multiple sympatric canids, there is resource partitioning of some kind - habitat, spatial, temporal, and/or dietary [Vanak & Gompper, 2009b; Kamler, et al., 2012]. However, dietary partitioning is often a consequence of spatial, temporal, or habitat partitioning [Vanak & Gompper, 2009b].

Since multi-species carnivore assemblages are seen almost globally, interference competition is now accepted as a crucial factor in the distribution and composition of mammalian communities [Estes et al., 2011; Ripple et al., 2014]. A number of studies have established that a “landscape of fear” exists not just in prey-predator interactions but even in intra-guild interactions. In the latter case, as is true for carnivores, dominance is typically based on size [Palomares and Caro, 1999]. This has given rise to the notion of “species-scapes”, which has been defined as a “spatial plane of species interactions that combines with resources and habitat structure to drive species distributions” [Fisher et al., 2013].

Conditions that allow several species to coexist could usually include either a minimum weight difference between species or massive character displacement, such as in the case of some parts of sub-Saharan Africa with the cape fox, bat-eared fox, and the black-backed jackal [Kamler, et al., 2012]. Within Indian ecosystems, the Indian fox (*Vulpes bengalensis*) has been shown to suffer from interference competition with the domestic dog (*Canis lupus familiaris*) [Vanak & Gompper, 2009a; Vanak & Gompper, 2010a]. A complete understanding of the habitat requirements for each species and how the species uses the landscape is an essential component of wildlife management. However, it is rare to find four species of canids occurring sympatrically. Despite the wide distribution of free-ranging dogs, it is also rare to find systems with multiple canids so that guild-level interactions can be understood.

The Banni grasslands of Kutch harbor five co-occurring canids: desert fox (*Vulpes vulpes pusilla*), Indian fox (*Vulpes bengalensis*), golden jackal (*Canis aureus*), and Indian wolf (*Canis lupus pallipes*) alongwith high densities of free-ranging domestic dog (FRD - *Canis lupus familiaris*). The Indian wolf however is rarely seen here, and may only be transient. Current understanding of the species ecology indicates that dogs are human commensals, jackals are likely to be habitat generalists, while the fox species are considered habitat specialists with the Indian fox tightly coupled with grasslands and the desert fox associated with arid areas (Figure 1).

**Figure 1.**
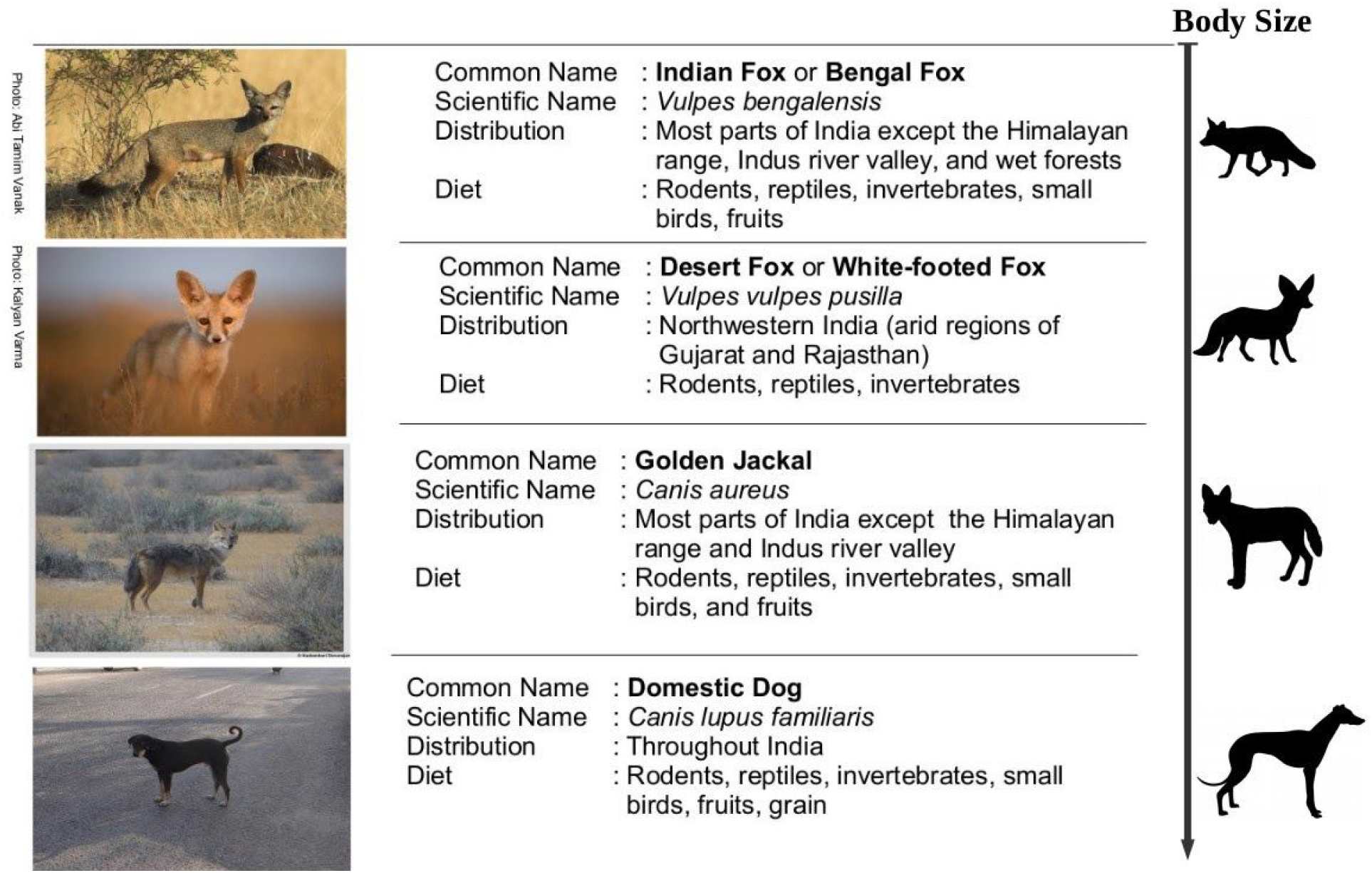
Brief natural history description of the four species of canids included in this study.

Here, we aim to understand how these multiple competing canid species interact over space and time, and the spatiotemporal associations facilitating their co-existence. We compare patterns of habitat use, as well as spatial and temporal segregation of four species of canids found in the study area (Figures 1 and 2).

**Figure 2.**
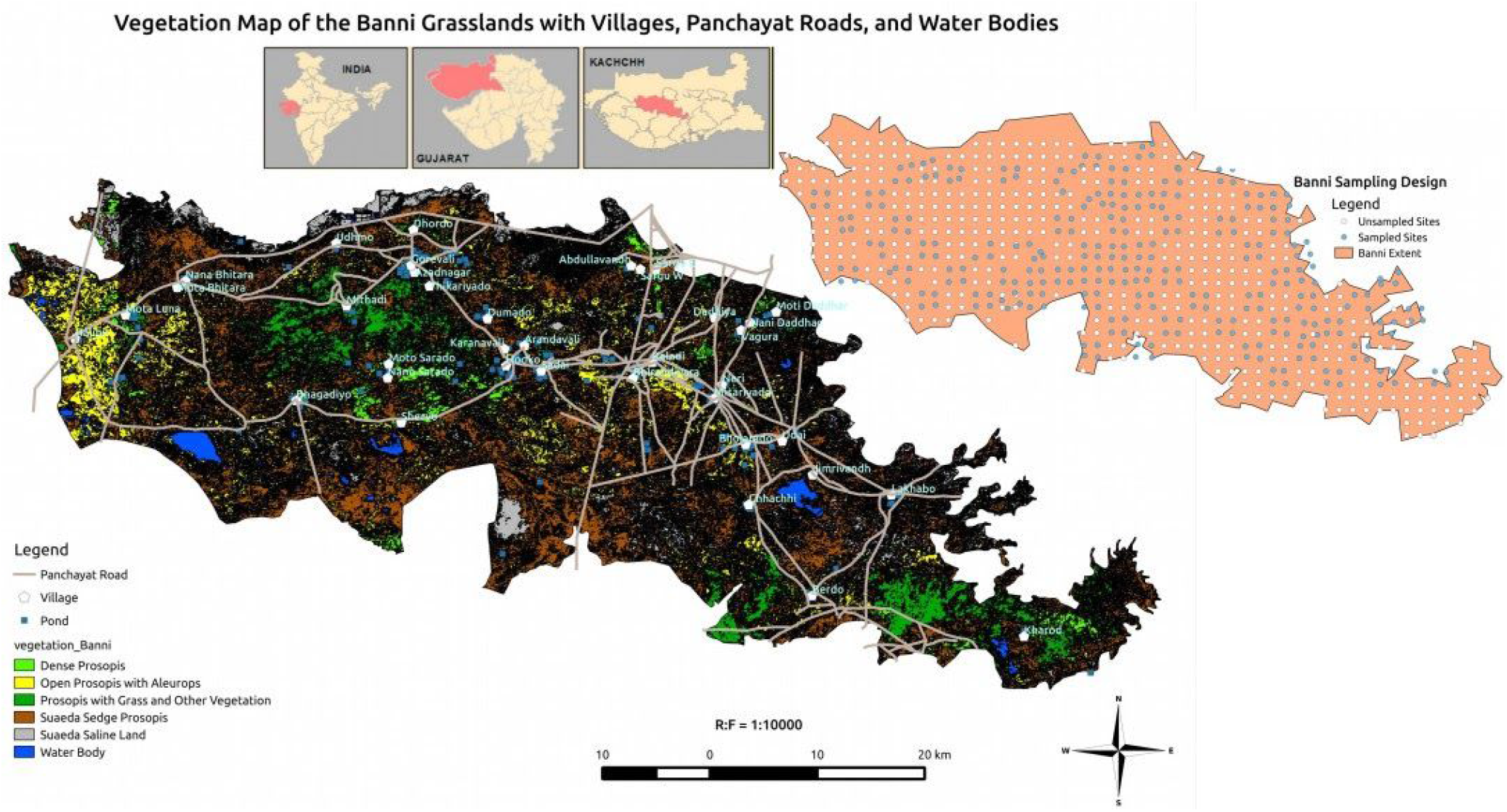
A map of the study area in the Banni grasslands depicting vegetation cover, roads, villages and water bodies along with some villages and adjoining regions, with the inset maps showing location of Banni in the Kutch district of Gujarat state in India as well as the sampling design employed. The gaps in sampling can be attributed to the presence of water bodies at some of the sites and due to an inability to obtain permission to set up camera traps in restricted defence areas. *Figure based on GIS layers courtesy of K-Link Foundation, India*.

We hypothesize that (i) the canids will vary in their distribution, with dogs and jackals more likely to be found close to human habitation such as villages, while the two fox species are less likely to be close to villages (Figure 4a), (ii) the presence of wild canids will likely vary based on habitat type with jackals widely distributed and found in areas with the invasive *Prosopis juliflora* (also called mesquite) and other mixed vegetation types, whereas the Indian fox is more likely to be associated with grassy areas, and the desert fox is more likely to be found in saline desert habitat (Figure 4a), and (iii) spatial overlap between two species will likely result in temporal partitioning or other behavioral character displacement between the species. We used a multi-species occupancy modeling (MSOM) framework to help account for imperfect detection and estimate the influence of the habitat and environmental variables, as well as the presence of the other species, on the co-occurrence of a given species (Figures 3–6).

**Figure 3.**
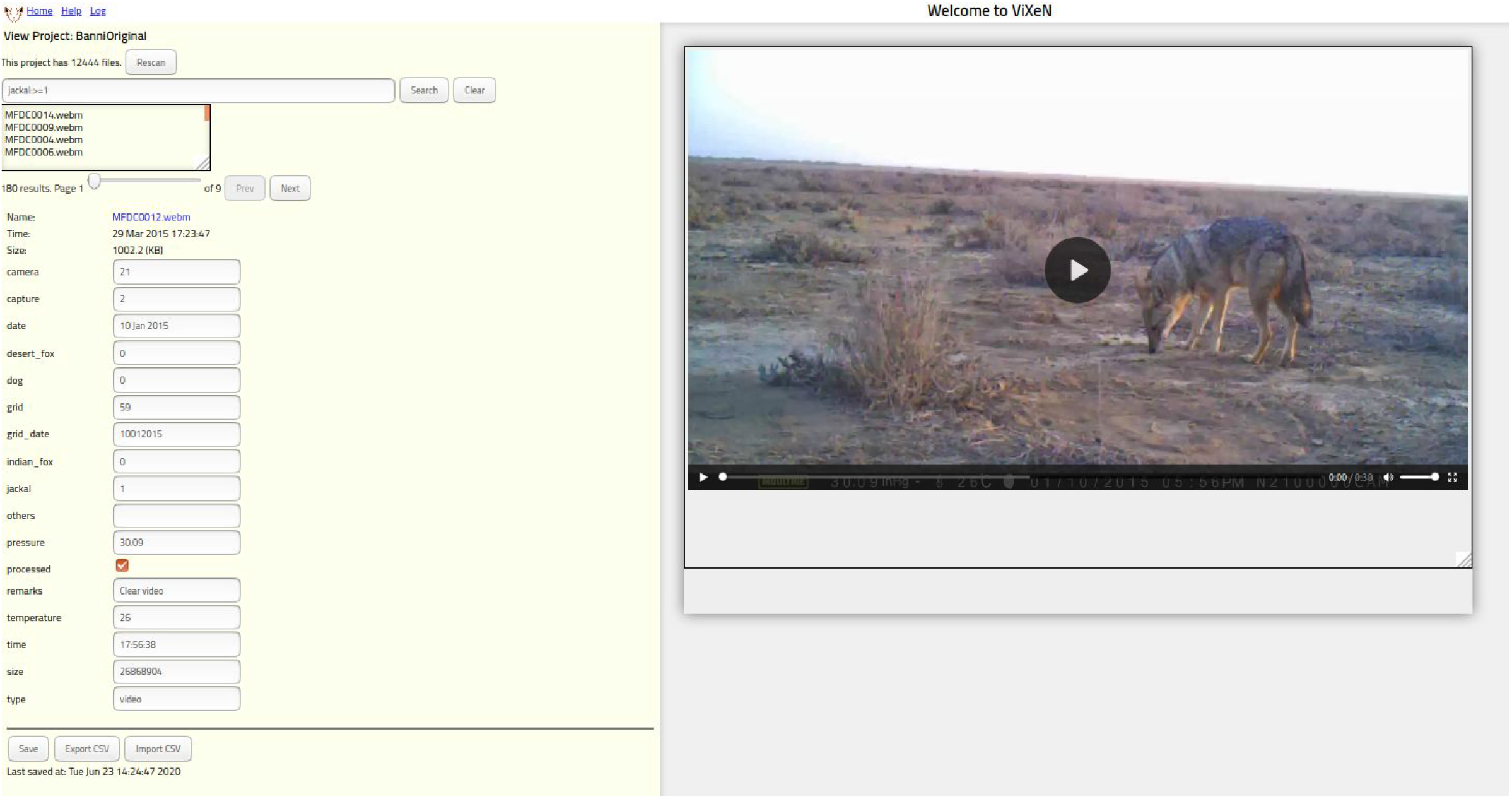
The ViXeN project interface for managing the camera trap videos with the corresponding tags that were created to extract variables of interest such as the species present in each video. The metadata seen to the left of the media viewer (used to view the camera trap videos converted to webm format) was extracted as a CSV file for subsequent analysis.

**Figure 4.**
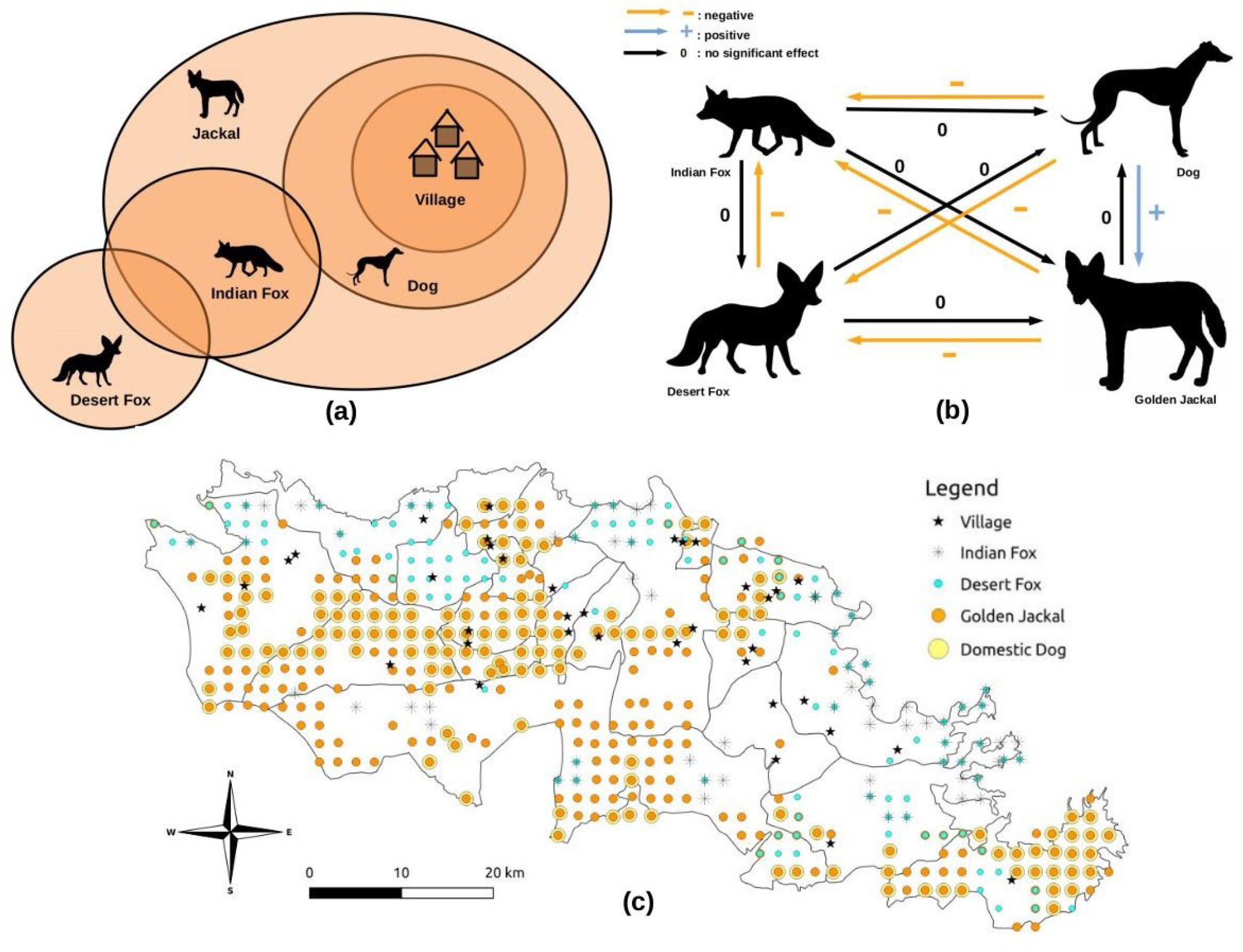
**(a)** This depicts the hypotheses the authors were testing which matches the distribution patterns for each species estimated using occupancy analysis as seen in (c). **(b)** The figure on the top right is a diagrammatic representation of the expected interactions between each species and the presence of the other three species in the area. The black arrows denoted by 0 indicate the body size-based assumption that the presence of this species has no effect on the species near the arrowhead. **(c)** A map of the study area in the Banni grasslands with the estimated occupancy (ψ) of the different canid species based on the highest probability of occurrence from the MSOM.

**Figure 5.**
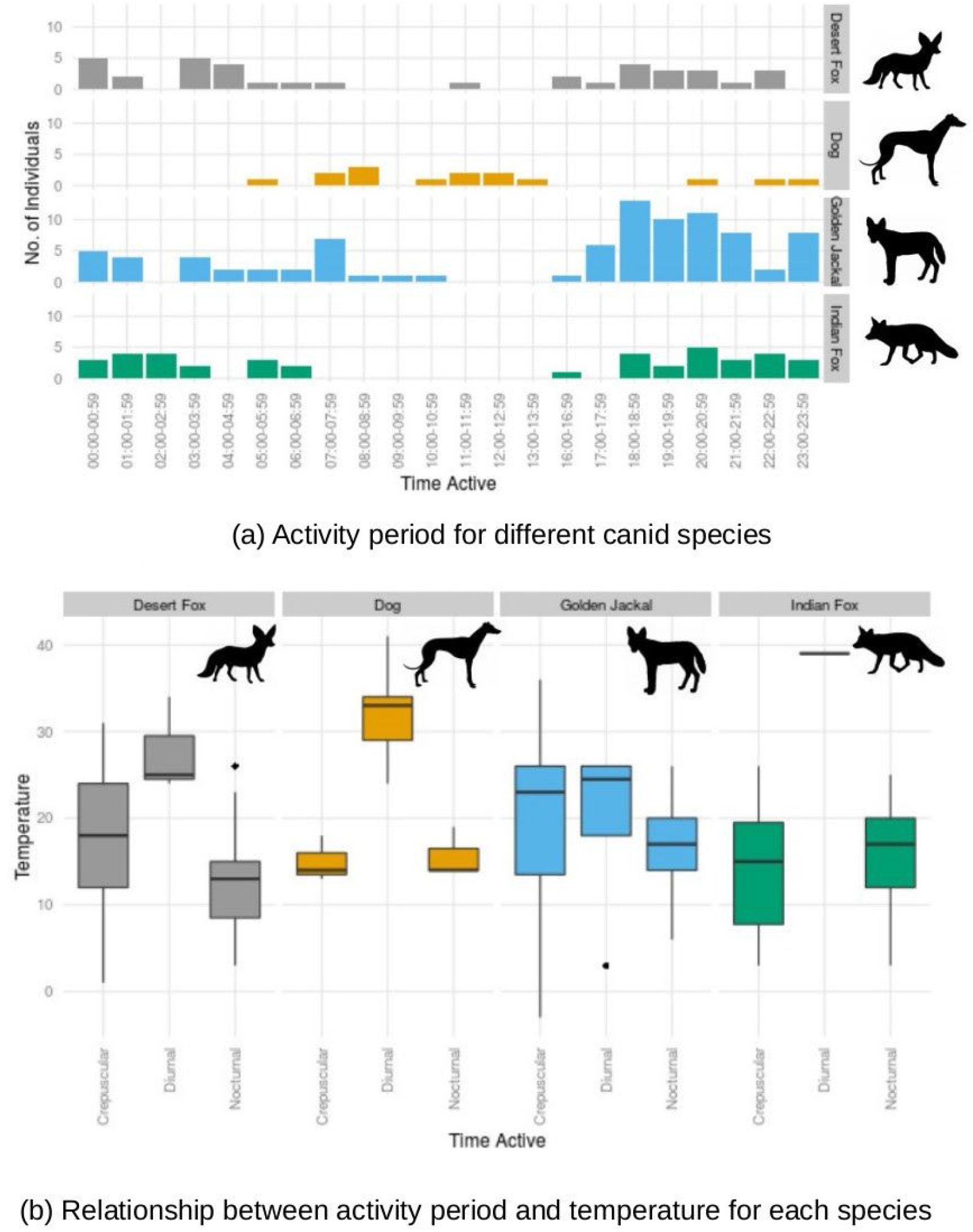
**(a)** Hourly activity count for each of the study species. The spatial overlap seen between the golden jackal and dog in the map in Figure 4 can potentially be explained by the temporal partitioning between the species seen here - dogs seem predominantly diurnal while the wild canids including the golden jackal seem to be crepuscular and nocturnal. **(b)** Boxplot comparing the relationship between activity periods and temperature for each of the study species. Temperature is in Celsius.

**Figure 6.**
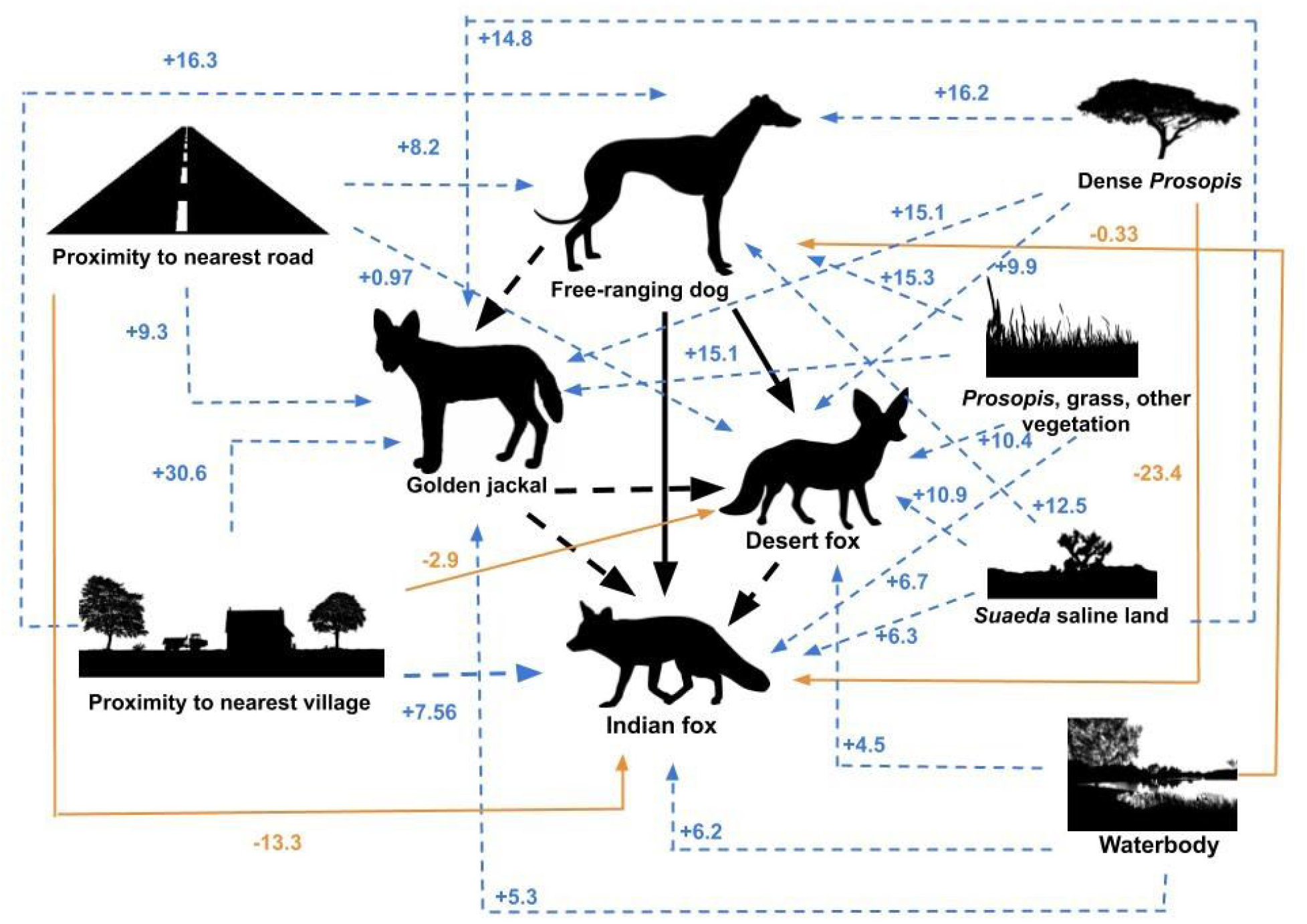
Visual summary of the interactions based on the MSOM using a body size-based hierarchy between the canids. For all colors, dashed lines represent a positive relationship while solid lines represent a negative relationship. The species-specific habitat covariate estimates obtained from the MSOM denote the degree and direction of influence of each covariate on the canids. The interactions between the individual canid species is obtained by comparing the conditional probabilities of co-occurrence using the interactions-specific estimates for each species based on whether other species are present or not present. Since dogs are the largest in the size-based hierarchy assumed, they influence all the other canids, although their occupancy is estimated using only the habitat covariates. Refer supplementary file shared through Figshare [Devarajan, 2020b] for model parameterization and model output used for this figure.

## Methods

### Study Area

This study was conducted in the Banni grasslands which lie in the Kutch district of Gujarat in north-west India (Figure 2) between September 2014 and March 2015. Spread over an area of 2500 km^2^, the Banni grasslands are considered the largest tropical grassland in Asia and the largest natural grassland in the Indian subcontinent. It is a mosaic of seasonal grassland patches and arid desert patches with salt pans.

The region has a high faunal diversity. Apart from the five canids, some of the other mammalian carnivores that can be found here include the desert cat, caracal, jungle cat, and striped hyena. While the Indian wolf has been an integral part of the carnivore assemblage historically, the species is very rarely seen in the region in recent years and hence was excluded from this study. Other species include over fifteen wild mammals with carnivores such as desert cat, jungle cat, and Indian grey mongoose, herbivores such as chinkara and nilgai, as well as Indian crested porcupine, long-eared hedgehog, black-naped hare, and Indian desert jird. Birds, reptiles, insects, livestock, and humans also emerged as bycatch from the camera trap study.

The spatial scale for this study reflects coverage of the whole Banni grassland landscape which is unique in that the canid community here comprises four co-occurring canids that are commonly found. The temporal scale was chosen on the basis of running a single season MSOM during the dry season, since it is logistically infeasible to set camera traps in the study area during the wet season when many parts of the area are inundated.

### Survey Design

Camera traps [Zielinski & Kucera, 1995; Silveira, et al., 2003; Sarmento, et al., 2010] were used to determine the occurrence of each of the species in order to understand habitat, spatial, and temporal partitioning between them. A grid-based sampling approach with a nested design was used by superimposing a 2 x 2 km grid over the entire study area of 2500 km^2^ in the Banni grasslands. We assumed that this cell size represented the home range of the largest species of wild canid (golden jackal) Aiyadurai & Jhala, 2006]. The home range of the Indian fox is between 1.6 and 3 km^2^ [Vanak & Gompper, 2007, 2010a, 2010b]. Since little is known about the ecology of the desert fox, we assumed a home range of about 4 km^2^ considering that it is intermediate in size relative to the jackal and Indian fox.

The cell size thus helps account for the assumptions of geographic closure and independence in MSOMs [Devarajan, et al., 2020]. The assumption of demographic closure is also satisfied since this is a single season study. While all four canids belong to the same guild and satisfy the crucial assumption of ecological similarity due to their relatedness within carnivores, they are easy to tell apart in the camera trap videos and hence satisfy the assumption of accurate identification, an important consideration in MSOMs [Devarajan, et al., 2020].

This grid-based chequerboard design for sampling resulted in 296 sample grids while a further 380 sites were taken as unsampled sites, giving a total of 670 sites for which the probability of occurrence estimates for each species were obtained (Figures 2 and 4c). We deployed a single Moultrie M990i No-Glow Game camera trap per grid for four consecutive nights as temporal replicates for modeling the detection probability, resulting in 1184 camera trap days in total.

Thirty two of these cameras were deployed in the field at any given time during the study. Since all cameras were from the same manufacturer, and belonged to the same model, any bias introduced from mixing camera trap types was avoided. In the absence of any sturdy trees on which camera traps can be securely mounted in the study area, custom camera trap mounts or stands were used.

In order to maximise detection if a species was present, a drop of lure (Cross Breed Food Lure from Kishel’s Scents, USA) was used for every camera trap. While this is considered an ‘active system’ method, lures are not as strong incentives as bait and hence unlikely to introduce any associated bias into the study [Garrote, et al., 2012; Gerber, et al., 2012]. For each camera trap site, the remotely-sensed covariates, such as the Banni extent, village locations, waterbody locations, roads, and vegetation (Figure 2), were obtained from land cover maps provided by the K-Link Foundation and Sahjeevan.

A body size-based hierarchy was assumed for the canids (Figures 1 and 4) and the interaction was built into the model for each corresponding species. In the model, this hierarchy was incorporated to account for occupancy estimates of a species for a site with and without one or more of the other species. Dogs are the largest among the canids in this study, followed by the jackals. The Indian fox is the smallest of the canids, while the desert fox is intermediate between the jackal and Indian fox in terms of body size. This assumption implies that the smaller canids are affected by interactions with the larger canids, while the larger canids are unaffected by interactions with the remaining canids. The Indian fox is thus influenced by the desert fox, golden jackal, and dog while the dog is not influenced by the other three canids.

### Analysis

The videos obtained from the camera trap were accessed using ViXeN [Ramachandran & Devarajan, 2018], an open source multimedia file manager for viewing the media, adding custom tags, and annotating metadata associated with media files such as videos, images, audio, and text (Figure 3). Custom tags representing the variables of interest were created. The associated variables for each video included information on whether any of the study species were present in the video and if so, the corresponding number of individuals (see Figure 3). These metadata of species occurrences were saved as a comma separated value (CSV) file. These metadata were combined with geospatial data based on the grid and camera trap numbers.

The resulting CSV data file was cleaned and exported for further statistical analysis and visualization [R Studio Team, 2014; Wickham, 2016; Wickham & Chang, 2016; Devarajan, 2020a] in R ver. 3.4.4 [R Core Team, 2013] and Python version 2.7.6 [Van Rossum, et al., 2007]. The Python libraries pandas [McKinney, 2015] and numpy [Oliphant, 2007; Virtanen, et al., 2020] were also used for the data cleaning and processing done in the analysis.

We extracted information on the ambient temperature at the time of detection and timestamp of canid captured in the camera, and used this metadata to infer a rough time-activity budget (Figure 5a) for each species. We used remote-sensing data to understand anthropogenic impacts on species-specific carnivore occurrences, as well as local site-level variation in habitat features.

### Multi-species Occupancy Modeling

The final occupancy estimate is represented by the species-specific probability *ψ*_ij_ and indicates the probability of site use across the landscape for each species (*i = 1,2,…, N*) at specific sites (*j = 1, 2,…, J*) and sampling occasions (*k = 1, 2,…, K*).

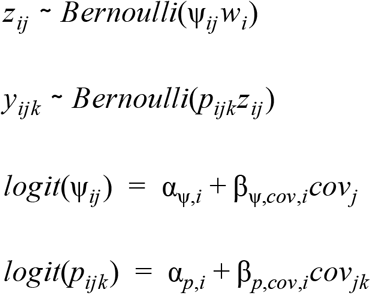

where *w_i_* is the indicator variable, Ω represents the probability that species *i* is part of the canid guild of size *N* = *4* in this case, is the true occupancy state matrix (if species *i* is present at site *j* then *z_ij_* = *1* else *z_ij_ =0*), α and β are linear predictors, and *y_ijk_* is the detection/non-detection of species *i* at site *j* over the *k*th sampling occasion having *p_ijk_* detection probability. The occupancy and detection probabilities (*ψ_ij_* and *p_ijk_* respectively) are modeled as a function of covariates (‘cov’) such as the proportion of dense *Prosopis* [DP] and proximity of each camera trap location (site) to the nearest village, simultaneously factoring in the occurrence of the larger canids based on the body-size hierarchy assumed.

The MSOM (see [Waddle, et al., 2010; Rota, et al., 2016; Devarajan, et al., 2020] for more information on MSOM implementation) was implemented under a Bayesian framework with JAGS using `jagsUI` in R. It was parallelized for a faster run using eight cores. The JAGS code provided as supplementary information through Figshare [Devarajan, 2020b] has the parametrization for each species. The model was parameterized with habitat covariates at the grid level (proportion in each grid of: dense *Prosopis* [DP]; *Prosopis*, grass, and other vegetation [PGOV]; *Suaeda* saline land [SSL]; water body [WB]) as well as proximity to nearest village and segment of road for each camera location along with the body-size based interaction described earlier. These were considered important in identifying the spatiotemporal patterns of distribution for all four canids. The estimates are based on three chains of 220000 iterations with burn-in of 6000 and adaptation of 12000 iterations, and a thin rate of 10, yielding 64200 samples from the joint posterior.

## Results

The camera trap survey yielded a total of 6221 videos of 30 seconds each from 296 camera locations for a total video footage duration of 3110 minutes. There were 315 videos with at least one canid identified and 797 videos with other organisms including livestock such as buffaloes, camels, horses, and goats (n=596), wild mammals such as other carnivores, herbivores, and rodents (*n*=142), birds (*n*=52), and invertebrates (*n*=5). The raw occurrences and co-occurrences based on the camera trap videos are provided in Tables 1 and 2. The spatio-temporal partitioning results are shown in Figures 4 and 5, while Figure 6 is a visual summary of the MSOM results. The model output for all species with the occupancy estimates and site-specific ψ estimates used to develop the map of the spatial interactions between the canids (Figure 4c) and the visual summary (Figure 6) is provided as supplementary information on Figshare [Devarajan 2020b] and includes the species-specific occupancy estimates, effects of interactions based on whether other species are present or not present, effects of habitat covariates on the occupancy of each canid species, and partial output of the site-specific occupancy.

**Table 1.**
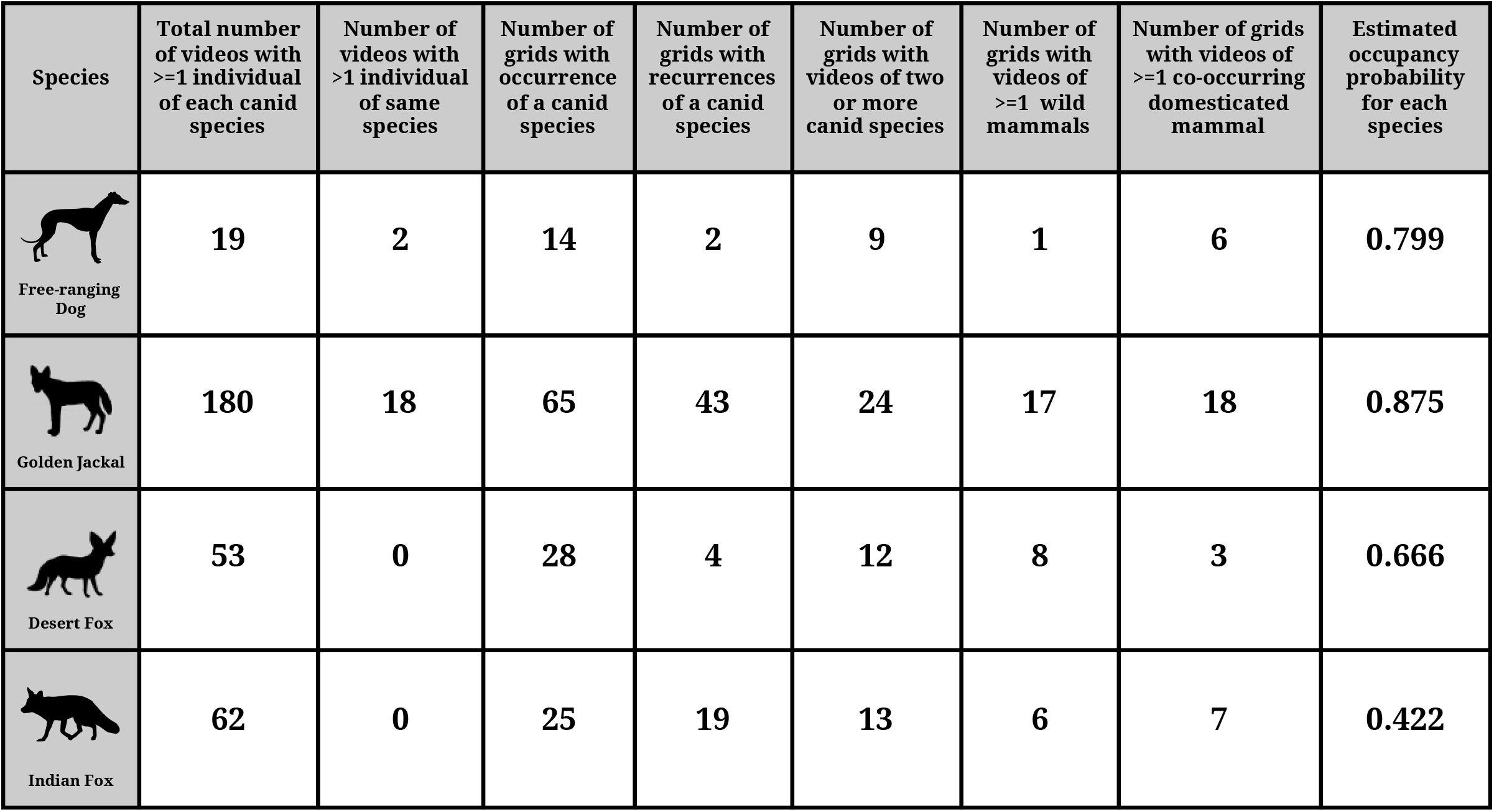
Site-specific species occurrences obtained from metadata obtained from camera trap videos annotated through ViXeN for the 296 sampled sites (camera locations) and 6221 videos based on the sampling design shown in Figure 2. The occupancy estimate (ψ) values shown for each species are from the MSOM using body size-based interactions and habitat covariates covering 670 (290 sampled and 380 unsampled) sites across the landscape as shown in Figure 2. Refer Figure 6 and the model output shared as supplementary information through Figshare [Devarajan, 2020b] for the species association estimates as well as the species-specific effects of the habitat covariates.

| Species | Total number of videos with $\geq 1$ individual of each canid species | Number of videos with $>1$ individual of same species | Number of grids with occurrence of a canid species | Number of grids with recurrences of a canid species | Number of grids with videos of two or more canid species | Number of grids with videos of $\geq 1$ wild mammals | Number of grids with videos of $\geq 1$ co-occurring domesticated mammal | Estimated occupancy probability for each species |
| --- | --- | --- | --- | --- | --- | --- | --- | --- |
| Free-ranging Dog | 19 | 2 | 14 | 2 | 9 | 1 | 6 | 0.799 |
| Golden Jackal | 180 | 18 | 65 | 43 | 24 | 17 | 18 | 0.875 |
| Desert Fox | 53 | 0 | 28 | 4 | 12 | 8 | 3 | 0.666 |
| Indian Fox | 62 | 0 | 25 | 19 | 13 | 6 | 7 | 0.422 |

**Table 2.**
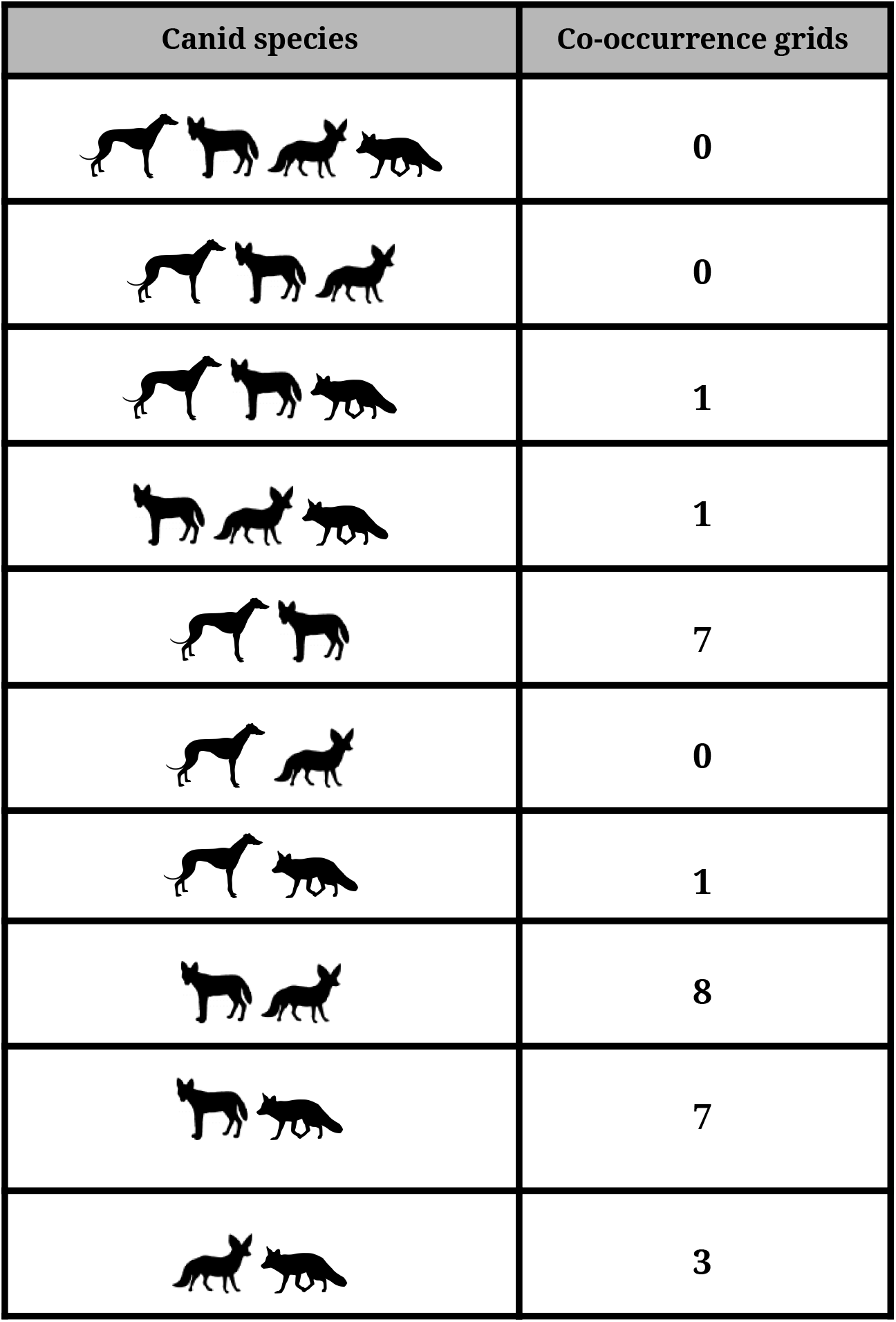
Site-specific species co-occurrences based on the camera trap videos obtained as described in Table 1.

We found a positive association between the golden jackal and dog, while the Indian fox shows avoidance of both of the larger canids (Figures 4 and 6). However, the desert fox was negatively associated with only dog presence. This is based on comparing the estimated conditional probability of occurrence for each canid species when one or more of the other species, on the basis of the body size hierarchy, was present or not present, and plotting the highest occupancy estimates of each canid on a map of the study area in order to glean patterns of spatial overlap in occurrence. The estimated occupancy is highest for the jackal and lowest for the Indian fox (Table 1).

We found that both the fox species were negatively associated with dog presence (Figure 6 and supplementary information on Figshare [Devarajan, 2020b]). There is some spatial partitioning between the Indian fox and the other canids, either directly or indirectly (Figures 4 and 6). Indian fox occurrence is also negatively associated with sites that had a higher proportion of *Prosopis*-dominated habitats (denoted by the covariate Dense *Prosopis*). They are also negatively influenced by proximity to the nearest road. However, while desert fox presence corresponded favorably with Indian fox occurrence, the former occur in more open *Suaeda fruticosa*-dominated saline habitats and are negatively influenced by proximity to human habitation such as villages.

The activity patterns for the canids based on the camera trap videos showed that free-ranging dogs were mostly diurnal while the wild canids were predominantly nocturnal or crepuscular. Both fox species were active at the same time, while none of the wild canids were active when FRDs were active. Jackals were crepuscular as well as nocturnal, while both foxes were mostly nocturnal.

When we combine the spatial patterns with the hourly activity data for the canids (Figures 4 and 5), we can see that although golden jackals and FRDs overlap spatially, they seem to be active at different times of the day, which indicates temporal partitioning between the species. On the other hand, while the two foxes seem to partition space, there appear to be pockets where they co-occur. However, there is a demarcation in habitat preferences between the two fox species - the desert fox is more likely to be found in saline and barren areas (*Suaeda* saline land) whereas the Indian fox is a known grassland specialist, with a preference for habitats with *Prosopis*, grass, and other vegetation (PGOV). Furthermore, the Indian fox is negatively associated with proximity to nearest road and areas with dense *Prosopis*, while the desert fox is negatively correlated with distance to nearest village (Figures 4 and 6). On the other hand, the golden jackal does not seem to be negatively associated with any of the covariates considered and has the largest occupancy estimate of all the canids.

## Discussion

In this study, we implemented a landscape-scale camera trapping effort, using a novel study design targeted at a guild of carnivores, with customized hierarchical modeling methods to estimate the occupancy of multiple species and extract the biotic and abiotic drivers influencing the distribution patterns of four co-occurring canids. We implemented a bespoke MSOM under a Bayesian framework accounting for imperfect detection, that takes into consideration the body size differences between the species. We coupled this with temporal data extracted from the camera trap study to understand the time-activity patterns of the canid species.

The results of our custom-fitted statistical modeling framework show complex patterns of spatiotemporal partitioning between the species. The larger canids, FRDs and golden jackals, are positively correlated with most of the covariates we considered and had the highest occupancy in the landscape. The distribution of the smaller canids, the desert fox and Indian fox, were negatively associated with several covariates, specifically the proportion of *Prosopis* in each camera trap grid and the proximity to the nearest village and road respectively. Furthermore, both fox species were negatively associated with dog presence. There emerged a strongly positive spatial association between FRDs and golden jackals, albeit with significant temporal partitioning between them. This suggests that spatial overlap between potentially competing carnivores at the local scale is often complemented with temporal segregation in order to facilitate coexistence at larger scales.

As visible in Figures 4–6, these results broadly match our hypotheses: (i) dogs and jackals were more likely to be found close to villages while the fox species were more likely to be negatively correlated with anthropogenic influences with the desert fox having a negative association with proximity to village and the Indian fox negatively correlated with proximity to road (Figures 4 and 6), (ii) canid occurrence did vary based on the habitat and time (Figures 4 and 5), and whereas jackals were positively impacted by the proportion of invasive *Prosopis juliflora* in the landscape, the Indian fox was negatively affected by this invasive (Figure 6), and (iii) the strong positive relationship between FRDs and jackals and resulting spatial overlap corresponded with temporal segregation between the species where dogs were almost entirely diurnal while jackals were nocturnal or crepuscular (Figures 4 and 5).

Recent research establishes that mere co-occurrence cannot be construed as evidence of ecological interactions [Blanchet, et al., 2020]. While the signal of interaction from observational studies is hard to establish, this study provides insights on how possible interactions as seen through the lens of spatiotemporal association and partitioning, along with habitat type and quality and human presence, affect the distribution and landscape use of a guild of sympatric carnivores [Karanth, et al., 2017]. Furthermore, there is evidence of potential interactions through repeated instances of different study species (and other small carnivores) captured on the same camera trap, often at different times of the day (see Figures 4, 5 and 6, and Tables 1 and 2).

The Banni grasslands, a seasonally-resource limited system, are rapidly being modified by the invasive *Prosopis juliflora* (commonly known as mesquite). This clearly has impacts on flora and fauna of the region. From habitat and dietary preferences of the canids, there emerges evidence that human-subsidized or commensal carnivores such as dogs and jackals are adapting better to this invasive species whereas the two fox species are negatively impacted. Furthermore, FRDs are considered invasives in many parts of the world including India [Home, et al., 2018] and are reservoirs of disease, and the rise in dog populations has been shown to have severe negative effects on wildlife including other carnivores as well as prey species [Home, et al., 2018]. Our research offers insights on how FRDs affect other carnivores within the same guild. The results of this study add to the body of evidence that indicates that the fox species, the smallest canids considered, are potentially negatively impacted by dog presence. This, coupled with the negative impact of mesquite and proximity to roads (Figure 6) in an area with increasing amounts of both (anecdotal evidence) as well as disappearing grassland habitats, is likely to adversely affect Indian fox populations, which already have the lowest occupancy of the canids in this study.

Globally, given that several carnivores are considered endangered, there has been an uptick in carnivore research and conservation projects. However, there are ecological, socio-economic, political, and epidemiological challenges to carnivore conservation. Possibly as a consequence of these challenges coupled with the resource-intensive nature of wildlife monitoring and the lag in statistical and computational tools and methods to handle ecological data, the conservation and management of carnivores has historically focused on single species. There is growing interest in studying entire communities simultaneously, paving the way for multi-species or community models in ecological research [Mackenzie, et al., 2002; Royle & Young, 2008; Royle, et al., 2009; Waddle, et al., 2010; Rota, et al., 2016; Sollmann, et al., 2011; Devarajan, et al., 2020]. An additional challenge to modeling communities is that co-occurrence alone is not sufficient to establish ecological interactions [Blanchet, et al., 2020]. Furthermore, incorporating species interactions into these models is considered challenging and hence remains an underutilized, if important, aspect of wildlife monitoring. Our study has utilized advances in modeling such as MSOMs combined with novel statistical analyses methods to account for imperfect detection and overcome some of these challenges.

In summary, through this study, we provide baseline information on the distribution of several carnivores. In addition, we explore intra-guild dynamics under an environmental gradient (using habitat covariates) in a human-dominated landscape (using proximity to villages and roads as proxies for anthropogenic influence). Furthermore, this research adds to the literature on a threatened and understudied landscape with multiple canid species. It also offers additional insights on how human-subsidized canids affect other canids in a community with multiple carnivores. The results of our study will hopefully prove beneficial in informing habitat and wildlife management at the community level, particularly in human-dominated landscapes that are vulnerable to precipitating changes from different angles.

## Acknowledgments

We thank the Gujarat State Forest Department for providing the permits necessary for this study. KD is grateful for the funding and institutional support received from the National Centre for Biological Sciences - Tata Institute of Fundamental Research (NCBS-TIFR), Wildlife Conservation Society (WCS) - India Program, Department of Science and Technology - Government of India, Thakar Jaikrishna Indraji Research Fund, Research and Monitoring in the Banni Landscape (RAMBLE) Project, and Sahjeevan. ATV was also supported in part through a grant from the National Research Foundation, South Africa (Grant No. 103659).

## Author Contributions

**Kadambari Devarajan:** Conception of research, planning research, data collection, data analysis, writing draft, reviewing manuscript. **Abi Tamim Vanak:** Conception of research, planning research, reviewing manuscript.

## Ethics Statement

The study was conducted inside and outside protected areas in Gujarat and research permits to carry out ecological research required for the study were obtained from the Office of the Chief Wildlife Warden - Gujarat (Permit No. WLP/28/C/150-52/2014-148) and the Gujarat Biodiversity Board. Since the methods used were non-invasive and protected species were not sampled, animal ethics committee approval was not required.

## Data Accessibility

Data used in this study can be accessed through Figshare: https://doi.org/10.6084/m9.figshare.13075097.v1 [Devarajan, 2020a]. The JAGS code to run the single season MSOM for all four species (assuming body size-based effects of interactions between the species) with six covariates as described in the Methods section along with the output obtained is also available through Figshare: https://doi.org/10.6084/m9.figshare.13089140.v1 [Devarajan, 2020b].

